## Supporting figure legends for "Image-based identification and isolation of micronucleated cells to dissect cellular consequences"

**Figure S1. Analysis of nuclear and MN characteristics after Mps1i addition to RPE1 RFP703/Dendra cells.**

**a)** Representative images of nucleus morphology classes in RFP703/Dendra cells after 24 h Mps1i treatment. The NLS-3xDendra2 channel (NLS-Dend.) is blown out to highlight the cytoplasmic signal. H2B = H2B-emiRFP703 channel. Arrows indicate chromatin bridges. Scale bar = 10 µm. **b)** Nuclei were manually scored for indicated phenotypes. Rare multinucleated cells were included in the misshapen nuclei class. Nuclei clearly in late telophase based on the cup shape of the chromatin were scored as normal, not misshapen. Msp1i- (DMSO): N = 2, n = 531, 833. Mps1i +: N = 3, n = 647, 597, 627 p: ** ≤ 0.01, *** ≤ 0.001, **** ≤ 0.0001 by Barnard’s test. **c)** Quantification of misshapen nuclei in cells manually labeled as MN+ and MN- from b). N = 3, n = 839, 1032. p: ** ≤ 0.01, by Barnard’s test. **d)** Histogram of number of MN per cell in Mps1i treated RFP703/Dendra cells. N = 5, n = 1323. **e)** Quantification of MN chromosome content in Mps1i treated RFP703/Dendra cells. Chromosomes enumerated using a CENPB PNA probe. N = 3, n = 86, 91, 82. Single chromosome MN are highly enriched in this population and MN containing chromatin fragments (0 foci) are rare.

**Figure S2.** **Description of VCS MN classifier training and assessment of MN assignment accuracy.**

**a)** Graphic depiction of pipeline for generating MN ground truth labels for VCS MN training and testing datasets. Two-channel images of Mps1i treated RPE1 RFP703/Dendra cells were acquired using 20x widefield epifluorescence. Nuclei were segmented on the H2B-emiRFP703 channel using the Deep Retina segmenter. These labels were used to generate 48x48 pixel crops centered on each nucleus. Crops underwent histogram normalization and filtering out of nuclei touching the image border. MN pixels were then manually annotated on each crop by an expert user. 2341 crops from 100 images (90 treated with Mps1i, 10 untreated) were annotated. **b)** Quantification of nucleus solidity in all nuclei versus nuclei labeled by the Deep Retina segmenter in RFP703/Dendra cells after Mps1i treatment. Highly lobulated nuclei (low solidity) were routinely missed by the Deep Retina segmenter. N = 1, n = 1357, 1492. p: **** ≤0.0001 by KS test. **c)** Graphic depiction of pipeline for training the VCS MN classifier. VCS MN was trained on the image crops and MN ground truth labels generated in (a). To accommodate Torchvision’s ResNet18’s expectation of a 3-channel 96x96 px image, each crop was resized and then expanded as follows: the first 2 channels are duplicates of the H2B-emiRFP703 image and the final channel is the result of Sobel edge detection. This image set was split into 3 parts for training, validation, and testing, as indicated, and fed into the VCS MN classifier. **d)** Representative images of MN assignment pipeline results. Automatic assignment depicted in top row (arrow) and manual assignment based on the NLS-3xDendra signal (NLS-Dend.) (arrow) in bottom row. Examples of correct and incorrect proximity-based assignment are shown. Scale bar = 10 µm. **e)** Quantification of proportion correctly assigned MN by proximity. Pooled proportion = 97%. N = 5 experiments, n = 1319 MN. **f)** Quantification of cell confluency in training images. Proportion of field covered by cells was defined by thresholding on the total NLS-3xDendra2 signal. Median = 33.2%. n = 30 images from 3 experiments. **g)** Distance between MN, nearest nucleus, and second nearest nucleus, centroid to centroid, in pixels. Median distances are 17.1 px (nearest) and 57.1 px (second nearest). n = 981 MN from images from 5 experiments. **h)** Distance between nuclei, border to border, in pixels in training images. Median = 30 px. N = 11 images, n = 507 nuclei.

**Figure S3. Analysis of VCS MN classification accuracy.**

**a)** Representative images of VCS MN analysis of RFP703/Dendra cells after Mps1i addition. Arrows on H2B-emiRFP703 channel indicate nuclear feature on left. Ground truth masks were annotated manually for nuclei (N) and micronuclei (MN). VCS masks and labels were generated automatically. In the VCS masks column, true positive MN are white, missed MN are green, and false positive MN are magenta. Nucleus blebs and chromatin bridges were rarely identified as MN by VCS MN. Instead, false positives were frequently associated with dim H2B signal (bottom). Examples of false negative MN show enrichment for large MN, MN overlapping the nucleus, and MN with dim H2B signal compared to nearby nuclei. Scale bars = 10 µm. **b)** Quantification of H2B-emiRFP703 intensity, MN area, and distance to nucleus for MN that were properly classified by VCS MN (true positives, TP), missed (false negatives, FN), or misclassified (false positives, FP). N = 5, 5, 1. n = 654, 327, 80. p: **** ≤0.0001 by KS test with Bonferroni multiple test correction. **c)** MN:nucleus NLS-3xDendra2 intensity ratios for manually classified intact and ruptured MN. Solid gray line = set VCS MN threshold. N=3, n=179, 113. **d, e)** Recall and rupture frequency for MN+ nuclei by # MN. MN+ cells were manually classified. N=2, n values on graph. Data show that nuclei associated with more than 1 MN are more likely to be correctly identified as MN+ by VCS MN (d) and that the presence of multiple MN increases the likelihood that at least 1 MN will be ruptured (e).

**Figure S4. Details of UNet architectures and object processing by the MNFinder module.**

**a)** The nucleus and MN (Nuc/MN) pixel classifier takes as input a cropped single channel image of chromatin and feeds it into two parallel, attention-gated UNets, one of which also has multiscale downsamplers (yellow). The nucleus weights from the basic UNet (top) are retained and MN weights from both UNets are fed to a third UNet for ensembling to produce the final MN predictions. **b)** Diagram of the triple decoder cell instance classifier. Two of the decoders have a UNet3+-like architecture with multiple skip connections and deep supervision during training. Feature depths are kept constant and most concatenation/max-pooling operations are replaced with addition to reduce training time. One decoder generates distance maps of a concave hull containing each nucleus and any associated MN (a “cell”) and the other generates a proximity map of each cell’s distance to all others. A third decoder uses a standard UNet with attention gates (cyan) to classify foreground pixels (nuclei and MN) and is used as input into every level of the distance- and proximity-map decoders via an integration block (magenta). **c)** Results from the nucleus/MN pixel classifier and cell instance classifier are further processed to improve accuracy. To limit misclassification of large MN as small nuclei, nuclei under a user-set area threshold are reclassified as MN. To limit MN undersegmentation, MN pixel groups are expanded by transforming each object to its convex hull. **d)** Distance and proximity maps from the cell instance classifier are combined to generate seeds for watershed segmentation. To correct for oversegmentation, only labels with boundaries that intersect a skeletonized version of the proximity map or background pixels are retained.

**Figure S5. Schematic of training data generation and use in MNFinder classifiers.**

**a)** Graphic depiction of pipeline to generate training data for the MNFinder MN and nucleus pixel classifier. A collection of single channel fluorescence images from several cell lines, chromatin labels, and microscope settings were manually annotated for nuclei and MN. Training, validating, and testing data were generated from these images by first normalizing image dimensions so that nuclei were approximately 30 px across their shortest diameter, generating random 128 by 128 px crops from all images, augmenting the data with a series of transformations, and then adding a second channel from a Sobel filter. This yielded a dataset of ~8800 images, which was split for training, validation, and testing as indicated. **b)** Graphic depiction of pipeline to generate training data for MNFinder cell instance classifier, where cell is a defined as a nucleus and its associated MN. The same starting image set was used as in (a). Ground truth associated nuclei and MN groups were manually labeled using the NLS-3xDendra2 cytoplasm signal. Grouped objects were then transformed into cells by drawing the convex hulls. These hulls were then transformed into distance and proximity maps using Euclidian distance formulas. These maps were combined with a mask of the foreground pixels to generate a 3-channel image. These images were randomly cropped and transformed to generate a training dataset of ~8800 images. This dataset was then split for training, validating, and testing of the cell instance classifier as indicated.

**Figure S6. Analysis of MNFinder errors.**

**a)** Representative images of MNFinder analysis from human fetal fibroblasts (DAPI, top), RPE1 RFP703/Dendra (H2B-emiRFP703, second) or U2OS RFP703/Dendra (H2B-emiRFP703, bottom two) cells. Arrows on the chromatin images point to the nuclear feature listed at left. Ground truth annotations for nuclei (N) and micronuclei (MN) were generated manually. Nucleus and MN masks were automatically defined using MNFinder. In the mask channel, true positive MN are white, missed MN are green, and false positive MN are blue. Neither nuclear blebs nor chromatin bridges were frequently miscategorized as MN. False negatives were enriched in small MN, and false positives were frequently nuclear lobes. Scale bars = 10 µm. **b)** Quantification of false negative (FN) MN area. MN missed by MNFinder were enriched in small MN across the testing dataset. TP = true positives, N = 12 images, n = 329, 60 MN. p: * ≤ 0.05 by KS test. **c)** Quantification of nucleus solidity in nuclei associated with false positive (FP) MN. False positives were frequently associated with highly lobulated nuclei with low solidity. N = 12, n = 329, 21. p: * ≤ 0.05 by KS test.

**Figure S7. Controls for VCS MN isolation experiments.**

**a)** Outline of RFP703/Dendra visual cell sorting validation experiment using CellTrace labeling as the activation trigger. Cells were incubated with CellTrace far-red and mixed with unlabeled cells at a 1:1 ratio. Nuclei were labeled based on CellTrace fluorescence intensity and converted with either an 800 ms (CellTrace+) or 200 ms (CellTrace-) UV pulse. The well was only partially converted prior to FACs analysis and sorting. Representative image of the mixed population prior to photoconversion is shown. Scale bar = 10 µm. **b)** FACS plot of Dendra2 red:green ratio versus CellTrace fluorescence. Colored bars represent gates. As expected, CellTrace+ cells generally have a high red:green ratio, CellTrace- cells a middle red:green ratio, and unanalyzed cells from both classes a low red:green ratio. Percentages on graph are the percentage of CellTrace- and CellTrace+ cells in their expected gate, i.e. the population purity. **c)** Histogram of CellTrace fluorescence in cells sorted on the Dendra2 red:green ratio in (b) after re-analysis by FACs. Values show a slight drop in cell purity for both populations after sorting. **d)** Data from Fig. 3B replotted with NLS-3xDendra2 red and green fluorescence intensity values plotted separately. Two peaks of red fluorescence intensity are visible immediately after photoconversion (0 h) and maintained over time. The increased population of nuclei at 4 and 8 h with only background levels of red fluorescence (Dendra2:Red <10^1.5^) likely represents unconverted nuclei moving into the imaging frame. The downward shift in red fluorescence intensity over time likely represents turnover of converted red fluorophores within the cells. N = 1, n = 82, 353, 285, 313. p: **** ≤0.0001 by KS test. **e)** Predicted classifier PPV (population purity) for low MN frequency U2OS cells (U2OS Broad). We observe a lower, but still substantial, enrichment of micronucleated cells in the MN+ population compared to a highly micronucleated population (e.g. Fig. 3C). N = 1, n = 17 cells.

**Figure S8. Differential UV pulses do not induce substantial transcriptional changes.**

**a)** PCA plot of cells treated with DMSO and exposed to 800 ms or 200 ms UV. **b)** MA plot of the data in a). Only 6 differentially expressed genes were identified in cells exposed to 800 ms vs 200 ms UV pulses and only 3 were downregulated over 1.5 fold: DDX39B, FASN, RGPD6.

**Figure S9. Analysis of visual cell sorting MN images.**

**a)** Manual analysis of nuclei features in cells labeled as MN+ or MN- by VCS MN in RNASeq experiments. Categories defined as in Fig. S1. N = 3, n = 875 MN+ nuclei and 1375 MN- nuclei. p: ns > 0.05 by Barnard’s test. **b)** Change in MN rupture frequency over time in asynchronous and synchronized cells treated with Cdk1i. Other = mitotic, MN-, or Dendra2- cells. N=1, n=~200 cells per time point. **c)** Change in MN rupture frequency between the start and end of a visual cell sorting experiment (4 h) and predicted change in classifier PPV due to ongoing rupture of intact MN based on values in (b).

**Figure S10. Controls related to Figure 6.**

**a)** Representative images and quantification of ATF3 nuclear mean fluorescence intensity in cells treated with DMSO or doxorubicin (Doxo.). N = 2 (colors on graph), n = on graph. **b)** Representative images and quantification of EGR1 nuclear mean fluorescence intensity in cells treated with DMSO or hEGF. N = 2 (colors on graph), n = on graph. For a-b: p: *** ≤ 0.001, by GEE. Scale bar = 20 µm. **c)** Same analysis as Fig. 6B, but with chromosomes in ruptured MN excluded from the foci count. Similar levels of aneuploidy were observed between groups as in Fig. 6B.
