## Supplementary material for "Image-based identification and isolation of micronucleated cells to dissect cellular consequences": List of supplementary tables

Table 1: Model testing. Comparison of NN model metrics for MN segmentation across cell lines

Table 2: MNFinder image dataset. List of all image set properties used for training, validating, and testing MNFinder.

Table 3: MN frequency. Quantification of MN frequency (percentage of cells with at least 1 MN) in images used to evaluate MNFinder.

Table 4: Comparison MN analyses. Describes the flexibility, sensitivity, specificity, and segmentation metrics of MNFinder compared to existing automated programs that quantify MN frequency in adherent cell images.

Table 5: DMSO 800 vs 200 DEG. List of differentially expressed genes (FDR ≤ 0.05) in DMSO treated RPE1 cells isolated after an 800 ms or 200 ms UV pulse. For all DEG analyses: padj = false-discovery rate adjusted p-values.

Table 6: Msp1i DEG. List of differentially expressed genes (FDR ≤ 0.05) in Mps1i versus DMSO RPE1 cells.

Table 7: Msp1i DEG log_2_FC. List of differentially expressed genes in Mps1i versus DMSO RPE1 cells filtered for log_2_ fold change above 0.58 or below -0.58.

Table 8: He et al DEG. List of differentially expressed genes (FDR ≤ 0.05) in RPE1 cells treated with nocodazole for 8 h versus control, initially reported in (He et al., 2018). Overlap with Mps1i DEG list noted in last column.

Table 9: Santaguida et al DEG. List of differentially expressed genes (FDR ≤ 0.05) in RPE1 cells that were treated with the Mps1i molecule reversine for 12 h, released, and ceased to divide, compared to control cells. Results initially reported in (Santaguida et al., 2017). Overlap with Mps1i DEG list noted in last column.

Table 10: Hallmark Mps1i. List of MSigDB Hallmark gene sets that are significantly enriched (FDR ≤ 0.05) in Mps1i treated RPE1 cells. For all Hallmark lists, NES = normalized enrichment score, padj = false discovery rate adjusted p-values, size = number of genes in gene set, leading_edge = , sig_genes_in_geneset = the gene names of set genes that were significantly upregulated.

Table 11: Hallmark He et al. List of MSigDB Hallmark gene sets that are significantly enriched in nocodazole treated RPE1 cells from (He et al., 2018).

Table 12: Hallmark Santaguida et al. List of MSigDB Hallmark gene sets that are significantly enriched in reversine treated RPE1 cells from (Santaguida et al., 2017).

Table 13: MN+ DEG. List of differentially expressed genes (FDR ≤ 0.05) in Mps1i treated RPE1 cells with MN versus without MN.

Table 14: MN+ DEG log2FC. List of differentially expressed genes in Mps1i treated RPE1 cells with MN versus without MN filtered for log_2_ fold change above 0.58 or below -0.58. Overlap with Mps1i DEG list noted in last column.

Table 15: Ruptured+ DEG. List of differentially expressed genes (FDR ≤ 0.05) in Mps1i treated micronucleated RPE1 cells with only intact MN versus at least 1 ruptured MN.

Table 16: Ruptured DEG+ log2FC. List of differentially expressed genes in Mps1i treated micronucleated RPE1 cells with only intact MN versus at least 1 ruptured MN MN filtered for absolute log_2_ fold change ≥ 0.58. Overlap with Mps1i DEG list noted in last column.

Table 17: Hallmark MN+. List of MSigDB Hallmark gene sets that are significantly enriched in Mps1i treated RPE1 cells with MN.

Table 18: Hallmark ruptured+. List of MSigDB Hallmark gene sets that are significantly enriched in Mps1i treated RPE1 cells with MN and at least 1 ruptured MN versus no ruptured MN.

Table 19: Log_2_FC per replicate. List of log_2_ fold change values for Mps1i DEGs with absolute log_2_FC ≥ 0.58 broken out by MN+ and rupture+ replicate. Genes with all NA values for either all MN+ or all rupture+ replicates were excluded from analysis. enriched_in_rupture+_cluster = genes present in cluster enriched in increased expression over Mps1i DEGs.
