## Supporting figures 1-10 for "Image-based identification and isolation of micronucleated cells to dissect cellular consequences"

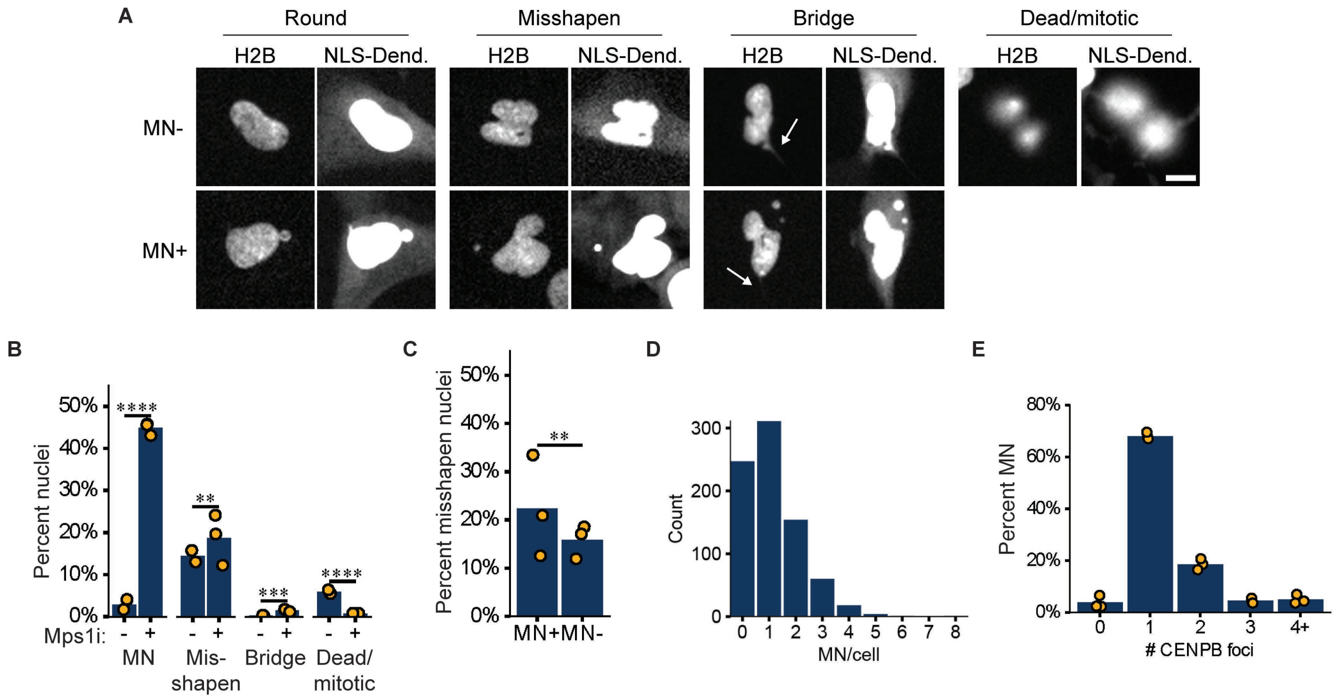


**Figure S1. DiPeso et al.**

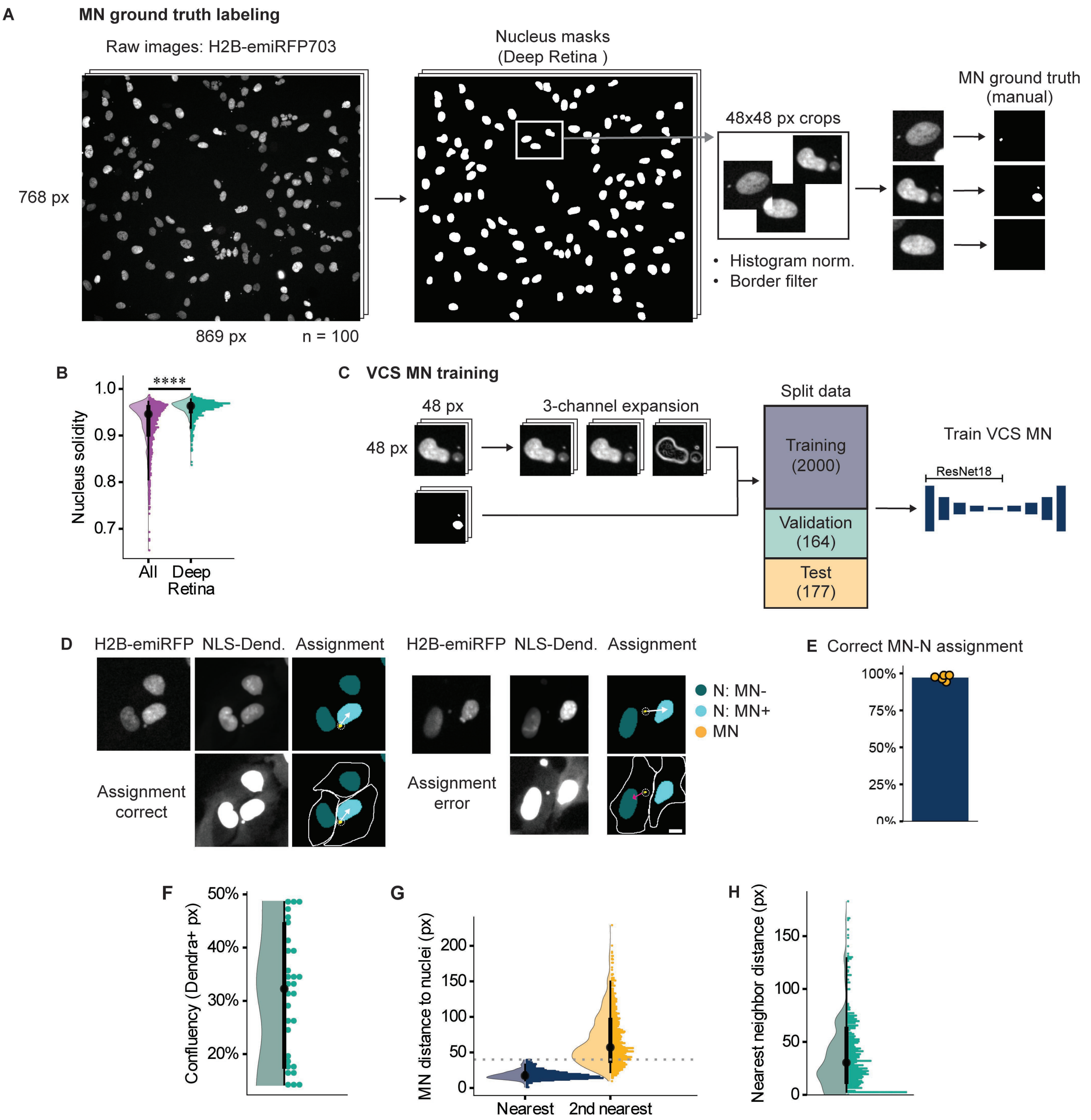


**Figure S2. DiPeso et al.**

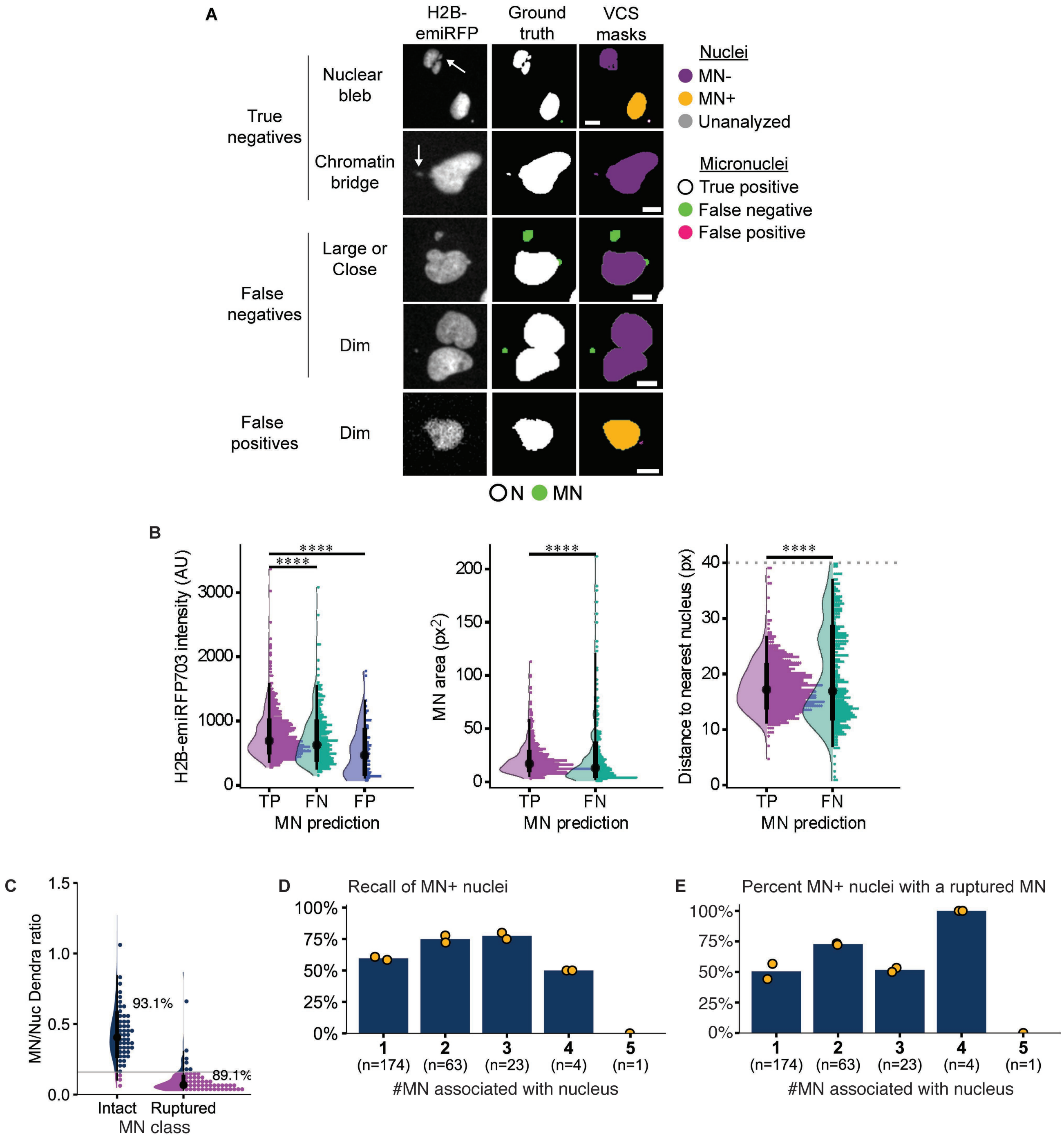


**Figure S3. DiPeso et al.**

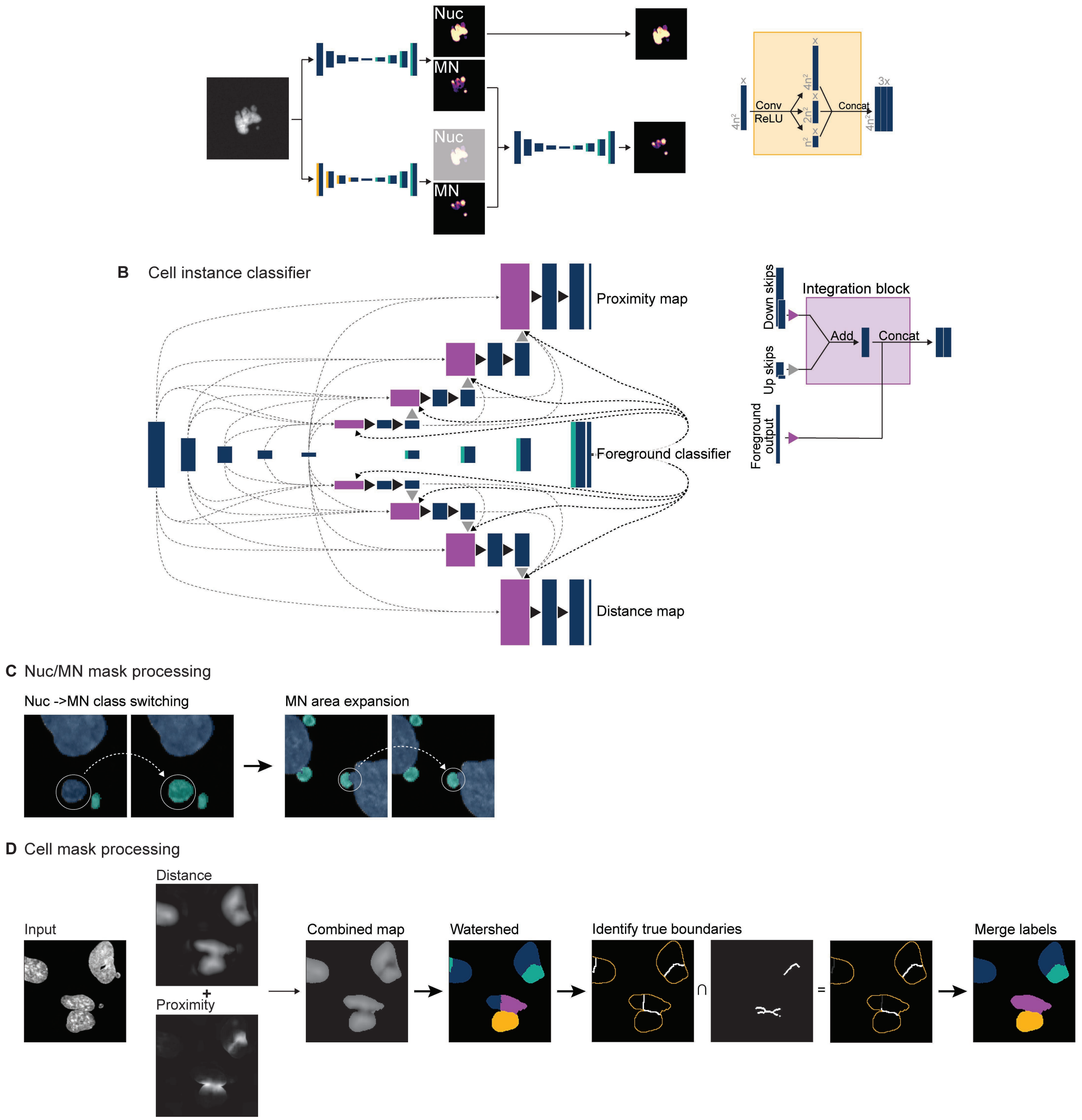


**Figure S4. DiPeso et al.**

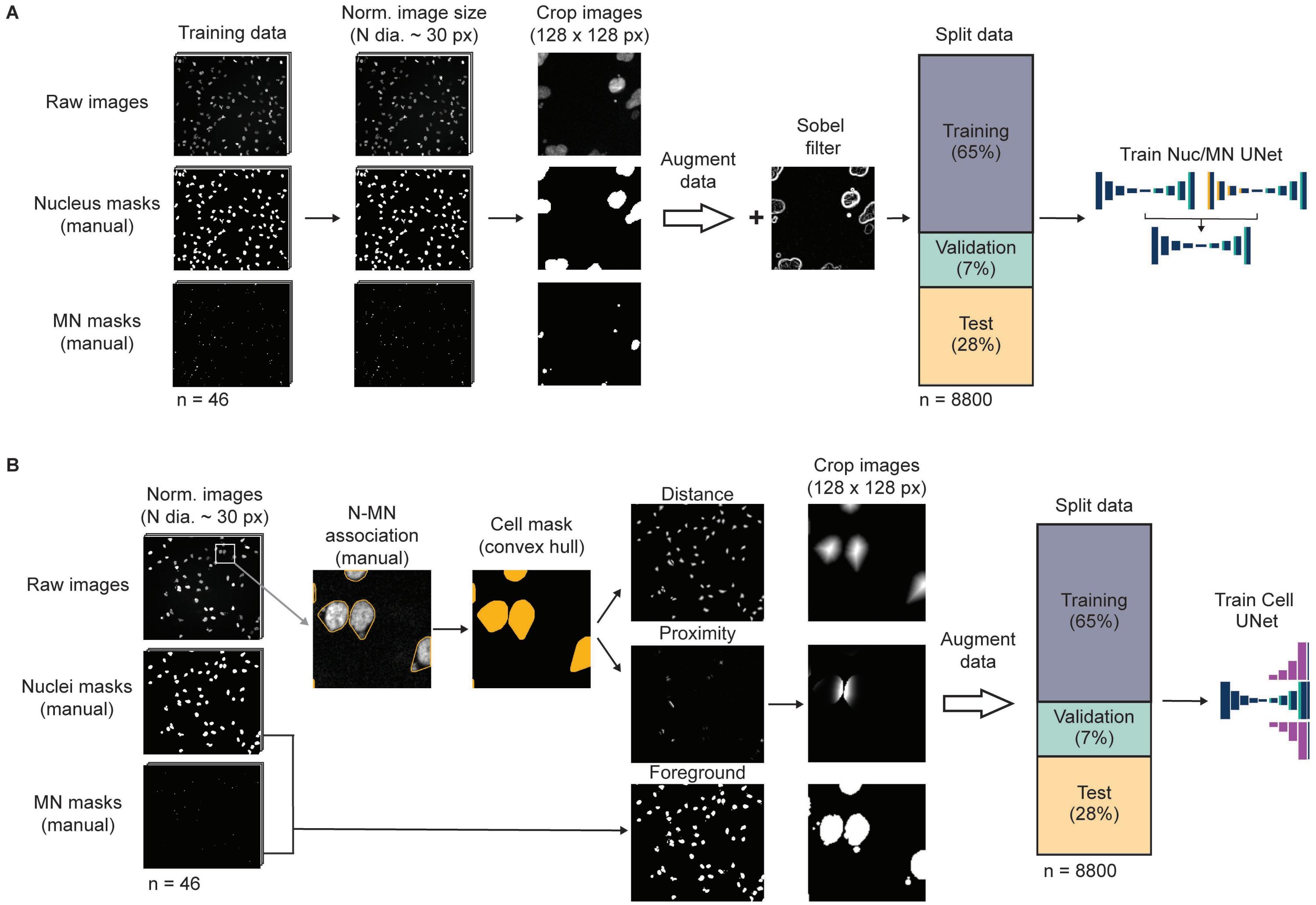


**Figure S5. DiPeso et al**

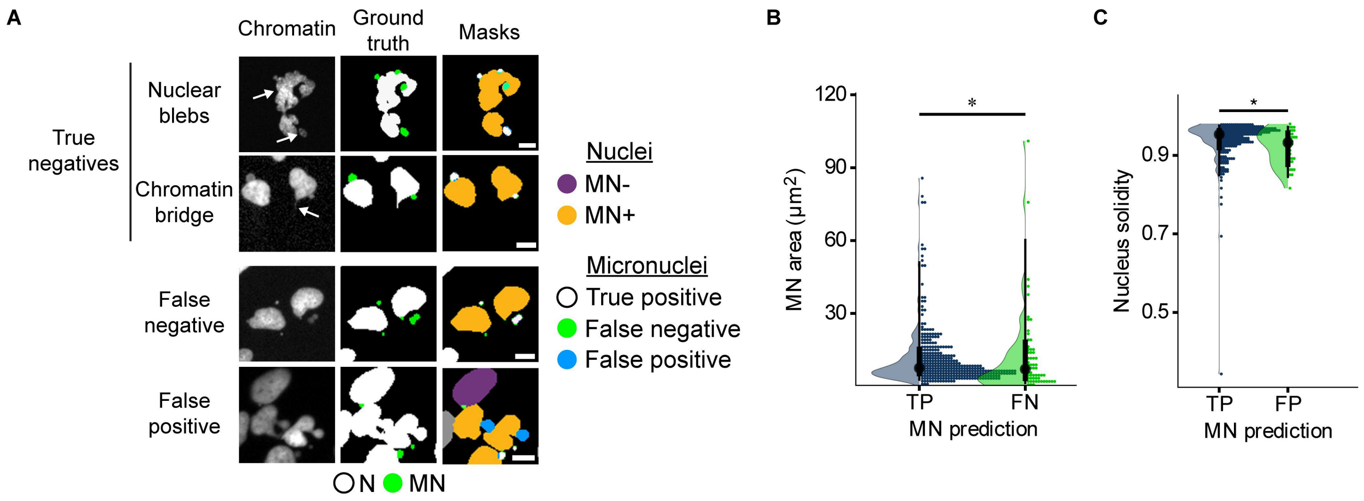


**Figure S6. DiPeso et al.**

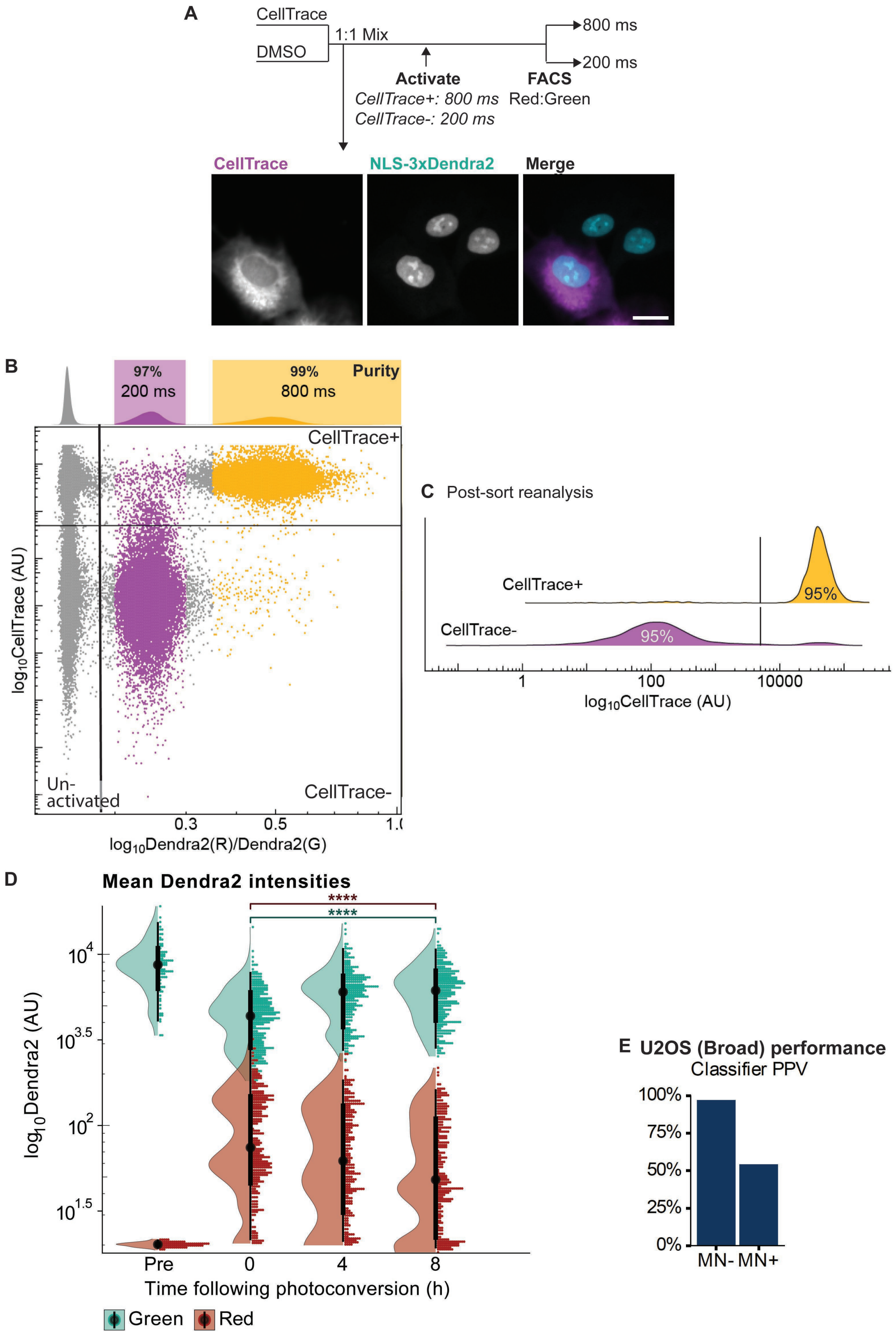


**Figure S7. DiPeso et al.**

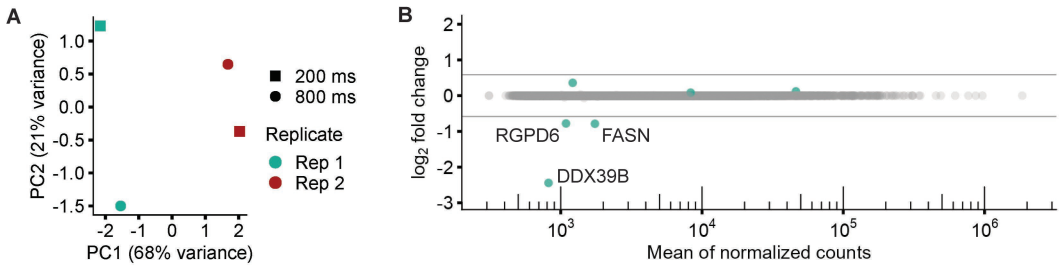


**Figure S8. DiPeso et al.**

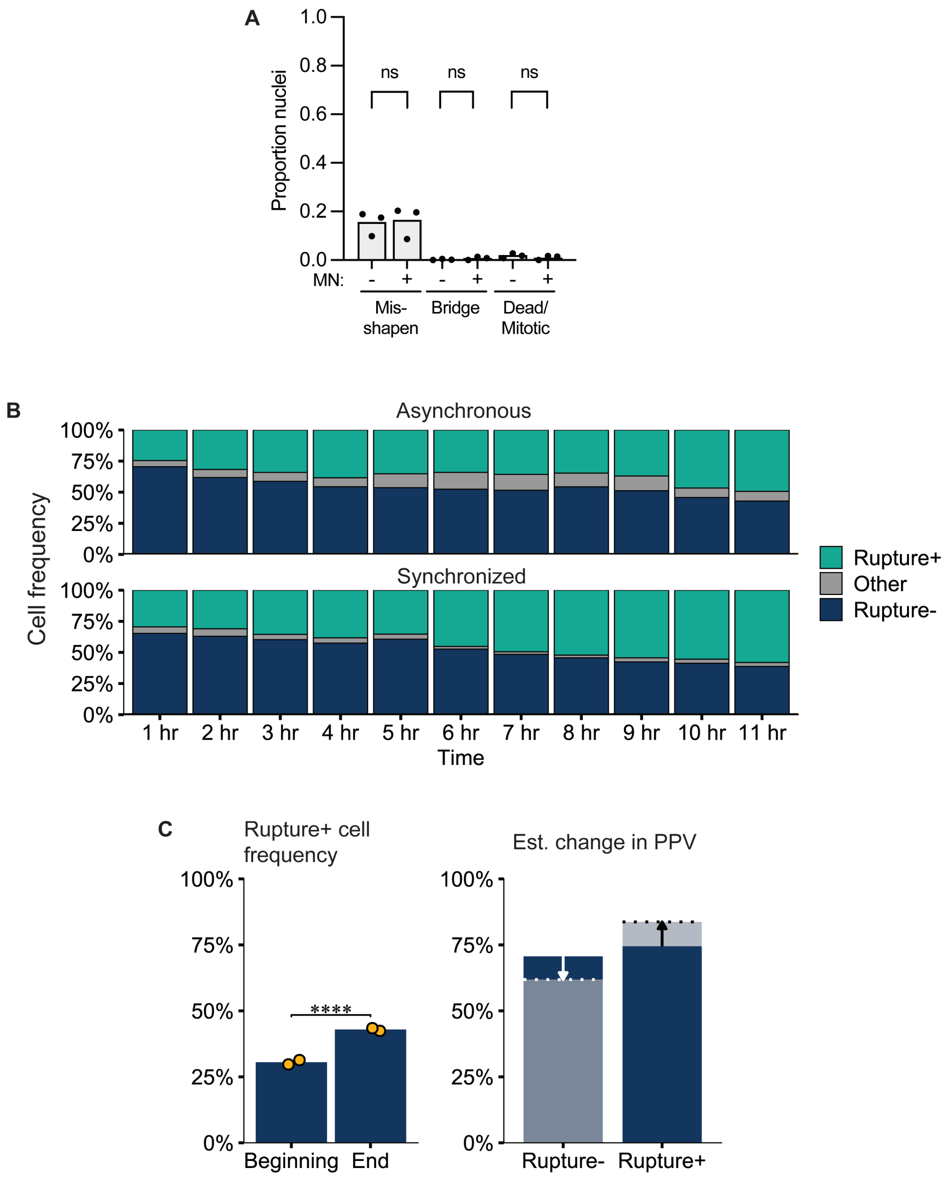


**Figure S9. DiPeso et al.**

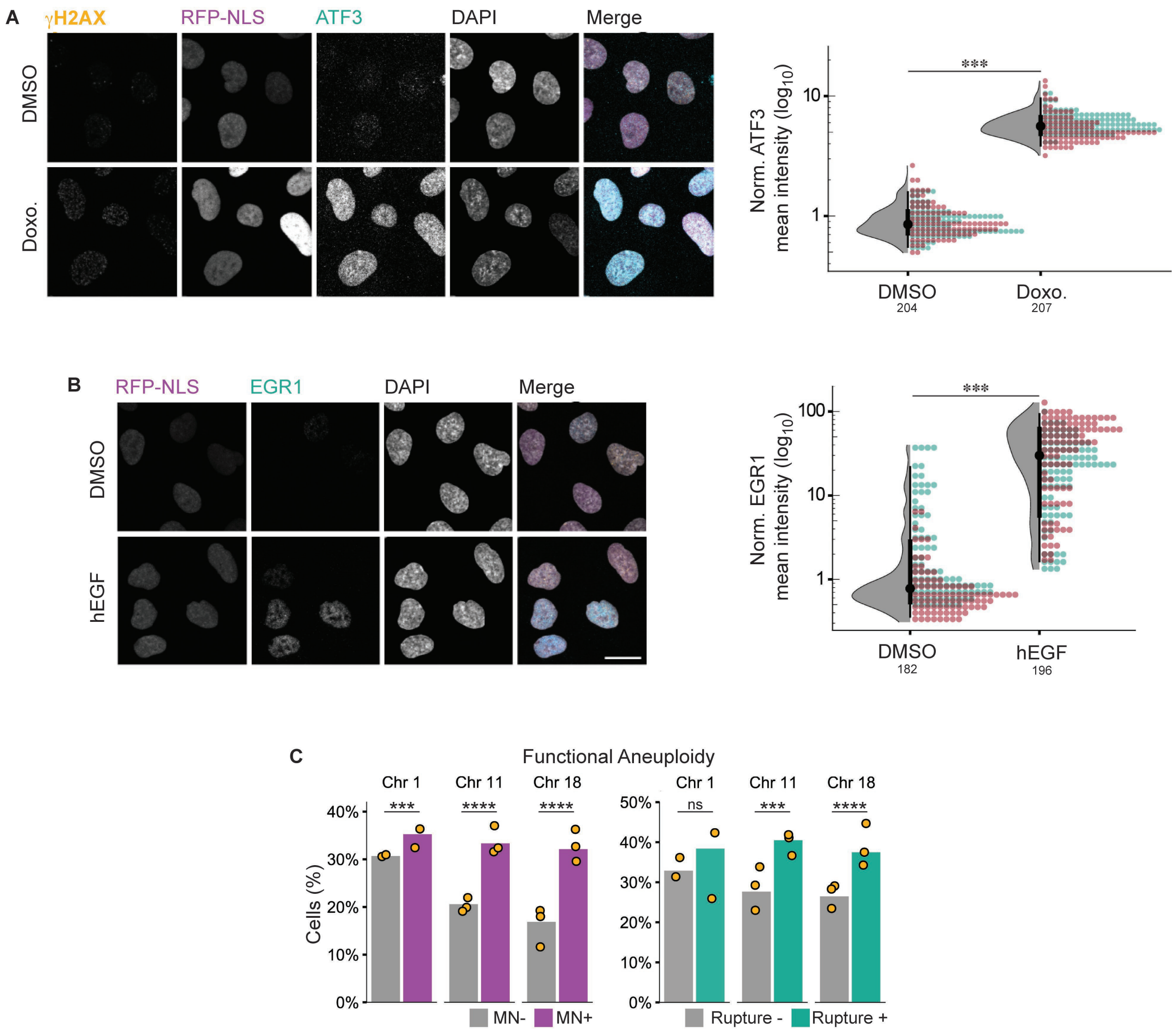


**Figure S10. DiPeso et al.**

-
-
